# Protein binding and differential distribution of calcitonin in the sub-compartments of blood: An In vitro study

**DOI:** 10.1101/2022.02.10.479976

**Authors:** Alex Bryan, Arun HS Kumar

**Affiliations:** School of Veterinary Medicine, University College Dublin, Belfield, Dublin-04, Ireland

**Keywords:** calcitonin, protein binding, compartmentalization, sustained effects, PKPD

## Abstract

The pharmacodynamics effects of therapeutically administered calcitonin is observed for extended duration much beyond its reported short half-life. The rationale for this pharmacodynamic-pharmacokinetics mismatch of calcitonin is not known. Hence in this study the hypothesis on sub-compartmentalization of calcitonin in blood to explain the extended pharmacodynamics effects of calcitonin was evaluated in vitro using the sheep blood model. A relatively higher proportion of calcitonin concentration was observed in the WBC compartment. The partition coefficient analysis showed levels of calcitonin to be higher on WBC membrane compared to intracellular. Molecular modelling to assess the binding of calcitonin with protein in plasma and WBC membranes indicated physiologically relevant higher affinity with beta-2-microglobulin’s in plasma and glycoproteins (CD44), sushi domains (CD25), fibronectins (CD206) and siglecs (CD22) on WBC membrane. The physiologically relevant higher affinity with several proteins on WBC membrane and plasma can be responsible for extended pharmacodynamics effects observed for therapeutically administered calcitonin.

## Introduction

Calcitonin (CT) is an endogenous regulator of serum calcium levels and is primarily produced by the parafollicular C cells in the thyroid gland.^1^ CT binds with the calcitonin receptor (CTR; present on osteoclasts) which enhances production of vitamin D and greater retention of calcium to increase the bone density. CT by activating CTR also becomes a potent inhibitor of bone resorption.^2-4^ CT is secreted in response to high plasma calcium levels and is vital to the physiology of calcium homeostasis.^1^ Hence synthetic CT is therapeutically used for conditions such as hypercalcemia, osteoporosis, osteoarthritis and Paget’s disease.^1,2^ Unlike the human CT (hCT), the amide bonds (b/w Lys-11 and Lys-18 amino acid) make salmon CT (sCT) more stable and absorbable from the GI tract with extended elimination half-life (54-76 minutes).^1,5^ sCT is well tolerated and has relatively less adverse effects.^2,5^ Although systemic administration of sCT is well tolerated among patients, its increased concentration in circulation can result in several undesired effects such as rhinitis, headaches, severe allergies and back pain.^6-8^ Hence novel formulations measures should consider this potential risk from increasing the concentration of CT in circulation.

Several formulations of CT have been developed for therapeutic use which have shown variable pharmacokinetics (PK) and pharmacodynamic (PD) effects.^9-11^ Using various formulations the bioavailability of CT has been improved up to 70% by parenteral delivery. Several studies have reported sustained pharmacodynamic effects (18-22 hrs) of CT which consistently seems to extend much beyond its relatively short half-life (40-60 mins).^9-11^ Recent study using nanomaterials and hydrogels have shown controlled release of CT up to 30 days with similarly extended pharmacodynamic effects in experimental model.^12^ Several attempts are also being made to enhance the pharmacokinetics of CT to further increase its bioavailability and achieve sustained delivery.^13,14^ The sustained pharmacodynamic effects of CT despite the limitations with its pharmacokinetics calls to question if such attempts to further improve the pharmacokinetics of CT are necessary especially when this can lead to several undesirable pharmacodynamic and adverse effects. Further the variability observed between the pharmacokinetics vs pharmacodynamic effects of CT may paraphs indicate a unique compartmentalization of CT in the blood.^15,16^ Such sub-compartmentalization of CT, if it exists, may be used to refine the therapeutic efficacy of currently available parenteral and oral formulations of CT rather than unnecessary efforts to newer CT formulations which carry a risk of increasing the CT concentration. Hence in this study we evaluated the sub-compartmentalization of CT with an in vitro experimental design using pharmacologically relevant CT concentrations.

## Materials and Methods

### Calcitonin Stock Preparation

Stock solution of salmon Calcitonin (sCT) was prepared by adding 1ml of DMSO (pH3.3) to 1mg sCT (Obtained from Polypeptide Group, CAS-no.: 47931-85-1) and vertexing it for 1 minute. A 10μl aliquot of this stock solution was added to 10ml of phosphate buffer saline (PBS) to get a working stock solution of sCT (10 μg/1ml). The working stock solution of sCT was vortexed and store in freezer until use.

### Blood sample and fractionation protocol

Intravenous blood samples were collected into heparin coated tubes from three healthy sheep’s at UCD Research Farm. The blood collection protocol was reviewed and approved by the Institutional Animal Research Ethics Committee (AREC-14-18-Kumar). 10ml of blood sample was transferred to 15 ml falcon tubes and 25μl of working sCT stock solution (calcitonin treated group) or PBS (control group) was added to the blood sample. The blood samples were then incubated at 37^0^C for one hour following which the blood samples were centrifuged (Rotanta 460/R centrifuge) at 2000xg for 15 minutes at room temperature (20^0^C). The plasma, red blood and white blood layers were separated from the centrifuged blood sample into separate tubes. Briefly the plasma was removed first and then the red blood cell (RBC) layer was removed carefully using a transfer pipette leaving about∼0.5 ml of the white blood (buffy coat) layer. Equal volume of Lymphoprep^™^ (Sigma-Aldrich) was added to the remaining buffy coat layer and centrifuged again at 2000xg for 15 minutes at room temperature (20^0^C) to isolate the white blood cell (WBC) sample. The isolated RBC component was similarly diluted with equal volume of PBS and centrifuged at 2000xg for 15 minutes at room temperature (20^0^C). The supernatant liquid (primary supernatant solution) from centrifugation of WBC or RBC component were also collected into separate tubes and were used for the assay of calcitonin levels using a commercially available ELISA kit (catalogue number S-1155.0001; Peninsula Laboratories).

### Cell Lysis Protocol

500μl of cell sample (RBC or WBC) was added to 1.5ml Eppendorf’s tube and centrifuged at 500xg for 5 minutes (Any supernatant was carefully transferred to the respective primary supernatant solution). 100μl of the cell pellet was resuspend in 300μl of celLytic^™^ MT reagent (Sigma-Aldrich) and incubated on a shaker for 15 minutes at 37^0^C. Following this the sample was resuspended in 300μl of PBS and was used for quantification of calcitonin levels using the ELISA kit.

### Quantification of calcitonin levels

Calcitonin was quantified using a commercially available Competitive ELISA kit. (catalogue number S-1155.0001; Peninsula Laboratories). All samples were stored at -80°C until used for analysis. Briefly, the reagents and samples were prepared as per the manufacturer’s instructions to establish the standard curve (figure 1) and quantification of calcitonin levels^17^. The levels of calcitonin in the samples following the cell lysate protocol described above was considered as the total intracellular calcitonin concentration in RBC or WBC. While the levels of calcitonin in the supernatant prior to cell lysis step was considered as the calcitonin concentration bound to cell surface (RBC-S or WBC-S). While the levels of calcitonin in the plasma was considered as total calcitonin (bound and unbound) concentration in the plasma. The calcitonin concentration is expressed as pg/ml and the calcitonin distribution is expressed in percent, with the total sum of calcitonin concentration in all five compartments [plasma, RBC, WBC, RBC-surface (RBC-S) and WBC-surface (WBC-S)] considered as 100 percent.

**Figure 1.**
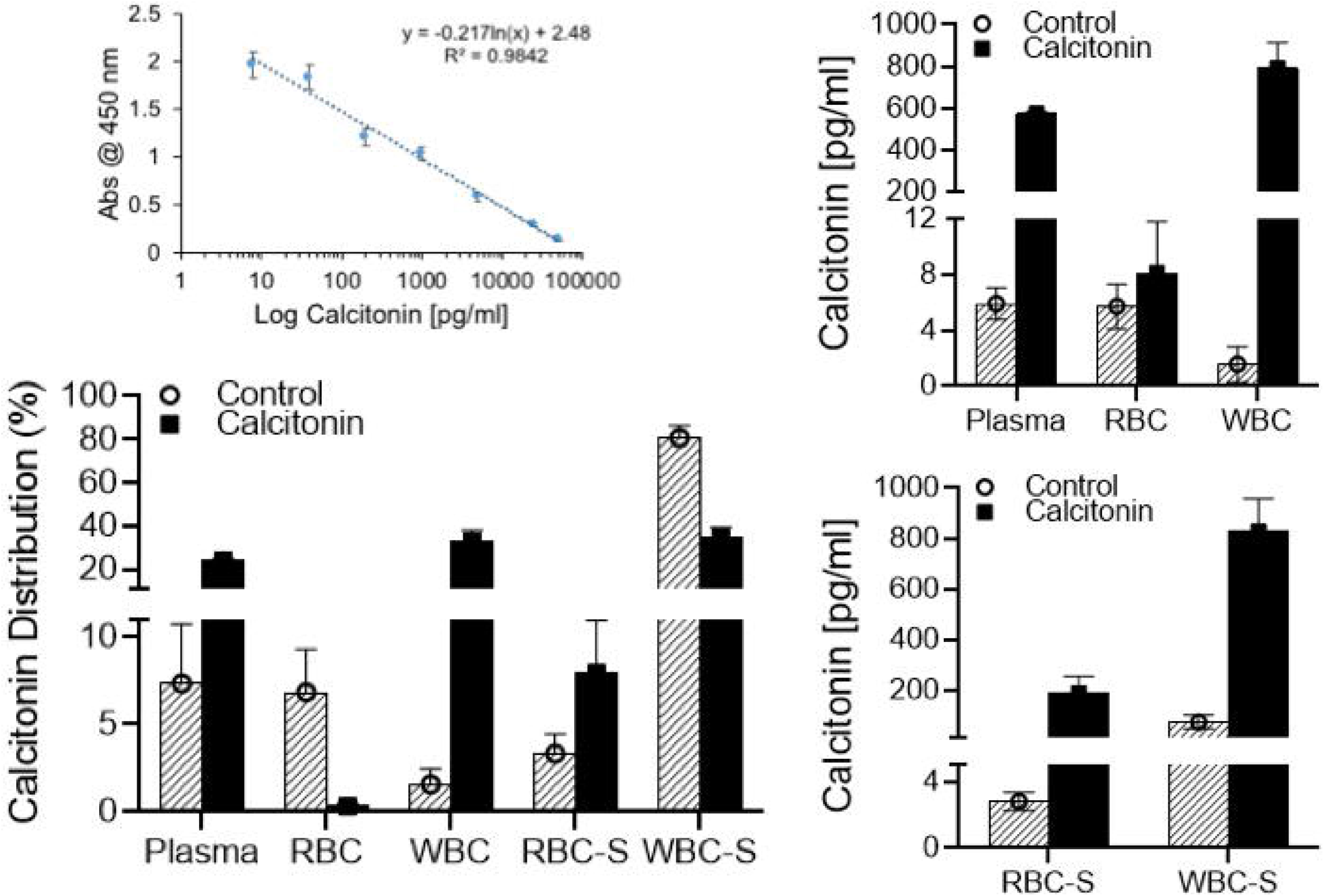
Distribution (percent) and concentration (pg/ml) of calcitonin in sub-compartments [plasma, RBC, WBC, RBC-Surface (RBC-S) and WBC-Surface (WBC-S)] of blood. The standard curve used for the estimation of calcitonin concentration is shown with the equation used inset in the graph. For the estimation of calcitonin distribution, the total concentration of calcitonin in all the five compartments was considered as hundred percent. The data are presented as Mean ± SD of three independent experiments in each group and each experiment was conducted in triplicates. The data between control and calcitonin treated group are statistically (p<0.05) significant.

### Partition coefficient of calcitonin in sub compartments of the blood

The partition coefficient of calcitonin was calculated by taking ratio of the calcitonin concentration in the respective compartments as reported previously.^18^

### Protein structure and molecular docking analysis

The protein data bank (PDB; https://www.rcsb.org/) was searched for major reported 3D structures of human plasma proteins and proteins on WBC membrane. 37 different plasma proteins and 30 different proteins on WBC membrane were identified and used for the analysis of molecular interactions (number of hydrogen bonds) with human-calcitonin (hCT) or salmon-calcitonin (sCT) using the Chimera software as reported before for other protein-protein interactions.^19-22^ The molecular interactions of sCT and hCT with the human calcitonin receptor ectodomain (PDB ID: 5II0) was considered as reference to quantify relative affinity of rest of the interactions. The reported EC_50_ values of sCT and hCT against human calcitonin receptor for cAMP production were used as reference value to generate the simulated dose repose curves as reported previously.^21,23^

### Statistical analysis

The data are presented as Mean ± SD of three independent experiments in each group and each experiment was conducted in triplicates. The data was analysed by two-way ANOVA using GraphPad Prism software (version 8).

## Results

In this study we evaluated the following five sub-compartments of the blood i.e., 1) plasma, 2) RBC, 3) WBC, 4) RBC-Surface (RBC-S) and 5) WBC-Surface (WBC-S). Differential distribution of calcitonin within these sub-compartments was observed (figure 1). In the control group, majority (∼ 82%) of calcitonin was located on WBC surface (WBC-S), with the rest of the calcitonin equally distributed between the plasma and RBC compartments (figure 1). Following exposure of whole blood to therapeutically achieved concentration (2.5 ng/ml) of calcitonin, its distribution within the blood compartments was significantly altered (figure 1). Following external addition of calcitonin, an equal distribution (∼ 30% each) of calcitonin in plasma, WBC and WBC-surface was observed with only less than 8% of calcitonin associated with RBC. The differences in concentration of calcitonin (pg/ml) in the sub-compartments (plasma, RBC, WBC, RBC-S and WBC-S) between the control (4.6-6.6; 4.6-7.6; 0.7-3; 2.2-3.3 and 46.1-99.6 respectively) and calcitonin treated (566.7-581.7; 7.3-11.7; 703.3-930.2; 138.4-262.9; 736.2-976.2 respectively) group were statistically (p<0.05) significant (figure 1). The total concentration of calcitonin in whole blood and plasma of the control group was 88±27 pg/ml and 6±1 pg/ml respectively. While in the calcitonin treated group the concentration of calcitonin in whole blood and plasma was 2400±73 pg/ml and 575±8 pg/ml respectively.

The partition coefficient between the different sub compartments of the blood was assessed (figure 2). The partition coefficient between the RBC-Plasma, WBC-Plasma, RBC-RBC-membrane(rm), WBC-WBC-membrane(wm) differed significantly (p<0.05) between the control and calcitonin treated group (figure 2). As the absolute concentration of calcitonin in the RBC compartment was very small, the partition coefficient difference in this compartment is of negligible clinical interest. However the variability in the partition coefficient of the WBC compartment indicated significant association of calcitonin with the WBC membrane. This was further evident from despite similar concentration (∼ 40%) of calcitonin both within WBC and on WBC membrane, a higher partition coefficient favouring calcitonin accumulation on the WBC membrane was observed in the calcitonin treated group. Among all the partition coefficients, the K_w/wm_ was most significantly influenced under higher calcitonin concentration, with the dynamics changing from intracellular accumulation of calcitonin at lower concentration (control group) to a marginal preference of cell surface association at higher concentration (calcitonin treated group).

**Figure 2.**
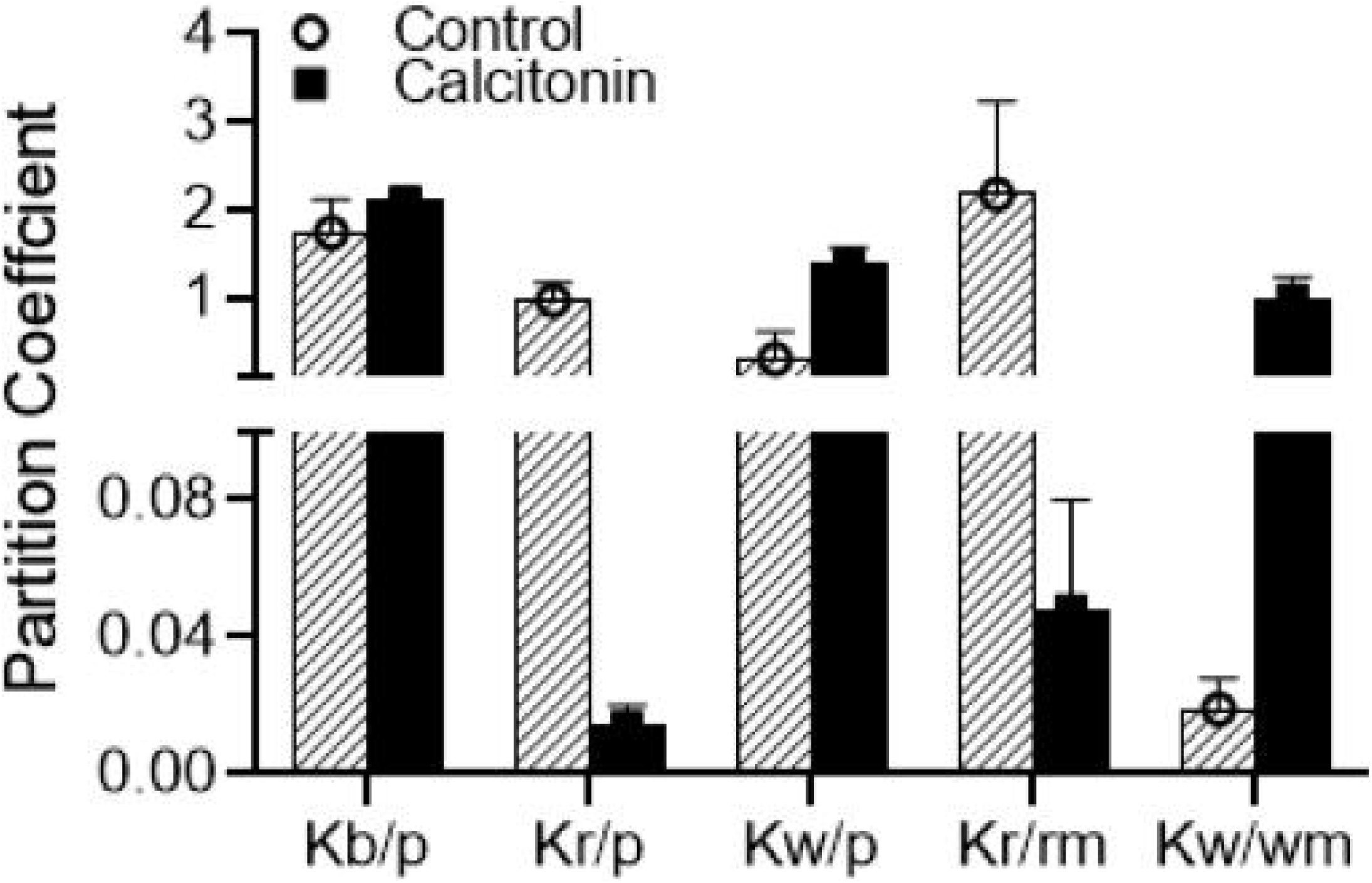
Partition coefficient (K) of calcitonin between various sub-compartments [blood (b), plasma (p), RBC (r), WBC (w), RBC-Surface (rm) and WBC-Surface (wm)]. The data are presented as Mean ± SD of three independent experiments in each group and each experiment was conducted in triplicates. The data between control and calcitonin treated group are statistically (p<0.05) significant for all categories except K_b/p_.

Calcitonin is reported to show reasonable plasma protein binding (∼ 40%). Hence to address the interaction of calcitonin with plasma protein or proteins on WBC-membrane, molecular docking analysis was performed to study the potential interaction of both salmon and human calcitonin with 37 major plasma proteins and 30 major proteins on WBC membrane (Figure 3A). Molecular docking analysis showed four fold higher affinity of salmon calcitonin against human calcitonin receptor ectodomain compared to the human calcitonin (Figure 3B,C, Table 1). hCT was observed to have higher affinity with the following proteins in plasma (3MRK (Alpha-Fetoprotein), 1DUZ (Beta-2-microglobulin)) and on WBC membrane [5VKJ (CD22), 6YIO (CD25)]. While the sCT showed significant affinity with the following proteins in plasma (1I4F (Beta-2-microglobulin), 1DUZ (Beta-2-microglobulin)) and on WBC membrane [4PZ3 (CD44), 5XTS (CD206)] (Figure 3D, Table 1). Both hCT and sCT were observed to significantly bind with several plasma as well as WBC membrane proteins with variable affinities (figure 3A). However sCT was observed to have superior affinity to WBC membrane proteins, while hCT in contrast showed superior affinity to plasma proteins (figure 3A).

**Table 1:** Affinity (EC_50_/Binding (nM)) of human or salmon calcitonin with human calcitonin receptor or selected proteins in plasma or WBC membrane.

| PDB ID | Human calcitonin | Salmon calcitonin |
| --- | --- | --- |
| *Receptor (5II0) | 7.2±1.3 | 1.4±1.2 |
| **Plasma proteins** |  |  |
| 1N5U | 6.61±1.19 | 2.10±0.60 |
| 3MRK | 1.21±0.22 | 11.33±3.24 |
| 1I4F | 2.31±0.42 | 1.31±0.38 |
| 1DUZ | 1.23±0.22 | 1.46±0.42 |
| 3PVN | 2.06±0.37 | 1.54±0.44 |
| 1EB1 | 2.05±0.37 | 2.25±0.64 |
| **WBC membrane proteins** |  |  |
| 5XTS | 8.05±1.45 | 1.73±0.49 |
| 1XIW | 12.43±2.24 | 1.98±0.57 |
| 1J8E | 8.63±1.56 | 2.22±0.64 |
| 4PZ3 | 3.05±0.55 | 1.30±0.37 |
| 2Z7X | 3.74±0.67 | 1710.8±488.8 |
| 6YIO | 2.98±0.54 | 1710.8±485.8 |
| 5VKJ | 2.66±0.48 | 1710.8±488.8 |

\* human calcitonin receptor ectodomain (PDB ID: 5II0)

**Figure 3.**
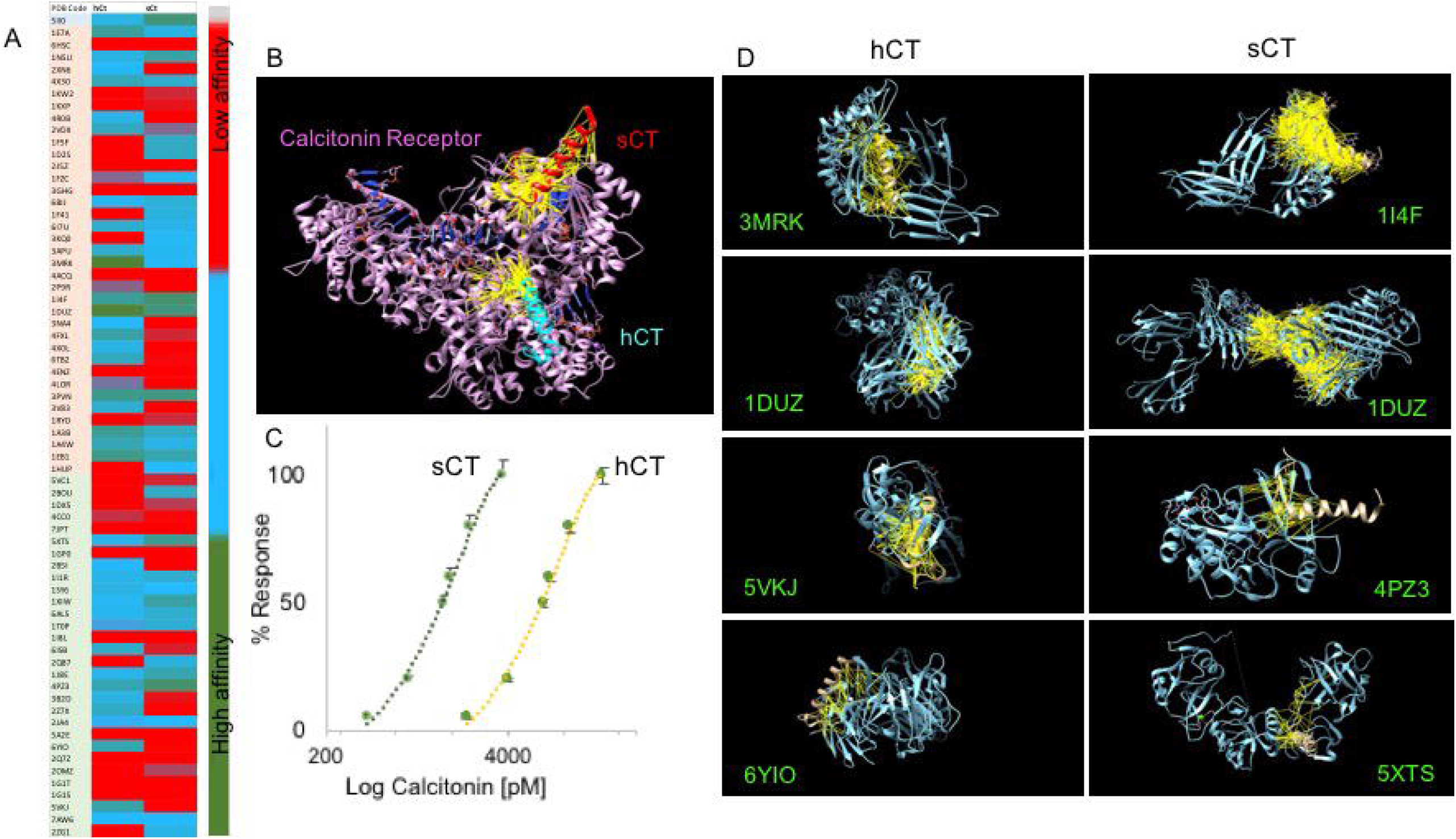
Molecular interaction of human (hCT) and salmon (sCT) calcitonin with human calcitonin receptor, various plasma and WBC membrane proteins. A) relative affinity of hCT and sCT with human calcitonin receptor (highlighted in light purple), various plasma (highlighted in orange) and WBC (highlighted in light green) membrane proteins. Scale: dark green represented strong affinity, sky blue represented medium affinity, red represented weak affinity. B) Molecular docking image of hCT and sCT with human calcitonin receptor ectodomain (PDB ID: 5II0). Yellow lines represent hydrogen bonds between calcitonin and its receptor. C) Simulated dose response curve of hCT and sCT based on its efficacy to produce cAMP levels following activation of calcitonin receptor. The data are presented as Mean ± SD of three simulation performed at ± 1, 2 and 3 sigma deviations. D) Representative images showing molecular interaction of hCT and sCT with selected plasma (3MRK, 1DUZ, 1I4F) and WBC (5VKJ, 6YIO, 4PZ3, 5XTS) membrane proteins. Yellow lines represent hydrogen bonds between calcitonin and its target protein.

## Discussion

Compartmentalization within the blood may be beneficial for less stable drugs, as protein binding in the sub-compartments will facilitate sustained pharmacodynamic effects of the drug.^24,25^ We report here such a behaviour for calcitonin, which in this study was observed to show selective affinity to the WBC sub-compartments both in the control as well as calcitonin treated groups. To the best of our knowledge such higher sub-compartmentalization of calcitonin in WBC is not previously reported. The specific sub-compartmentalization of calcitonin in WBC and plasma may paraphs explain the extended duration of pharmacodynamic effects of therapeutically administered calcitonin despite its short half-life. The observations from this study has several implications to both development of novel calcitonin formulations and as well as clinical use of currently available calcitonin formulations.^5,9,11,12^

Blood consists of cellular (RBC, WBC and platelets) and fluid (Plasma) compartments and drugs have equal opportunity to be uniformly diffused into all the sub-compartments.^26^ However selective molecular interactions of the drug will preferentially facilitate presence of the drug in specific sub-compartments.^27,28^ The molecular interactions could be with specific proteins, other biomolecules, selective transporters or channels on cell membrane or in plasma. In this study we observed that calcitonin was predominantly (82%) present in the WBC surface in the control group. The calcitonin concentration on WBC surface was∼ 15 times higher than that observed in the plasma in the control group. This observation is of interest in interpreting the literature on calcitonin pharmacokinetics as most studies in humans have reported a calcitonin concentration of 5-15 pg/ml in plasma or serum,^29,30^ which based on observation from this study may represent <8% of total calcitonin concentration in the whole blood. Hence the reported values of calcitonin concentration in plasma or serum in our opinion should be readdressed considering the potential of calcitonin to sub-compartmentalization in WBC.

The plasma concentration of calcitonin (6±1 pg/ml) observed in the control group in this study was far lower than that reported in sheep (165.13±12.53 pg/ml) by other studies.^31-33^ However these previous studies have used less sensitive/specific calcitonin assays.^32,33^ Also it is not known if breed specific differences can contribute to the variance observed in calcitonin concentrations. Another likely explanation for observing lower concentration of calcitonin in this study could be poor sensitivity of our assay against sheep specific calcitonin in contrast to its higher sensitivity against salmon or human calcitonin.^17^ Hence the absolute values of calcitonin levels reported by this study in the control group should be interpreted with caution. However this potential sensitivity issue should not affect our interpretations of relative distribution of calcitonin between the various sub-compartments.

The influence of drug dose on its pharmacokinetic parameters is widely reported In the literature.^34^ Hence the observation of significant changes to the sub-compartmentalization of calcitonin at a higher but therapeutically relevant concentration although previously unreported is not surprising. Even at the therapeutically relevant concentration of calcitonin the major sub-compartments of relevance were plasma and WBC, as the concentration of calcitonin in the RBC compartment was very low to be pharmacodynamically significant. Although this study didn’t analyse the calcitonin levels in the platelet compartment, but accounting the total calcitonin added with levels of calcitonin in different sub-compartments evaluated, in our opinion the platelet compartment is an insignificant entity for influencing pharmacodynamic effects of calcitonin. Between the plasma and WBC, this study observed a marginally higher presence of calcitonin in the WBC compartment at therapeutic concentration. This preferential presence of calcitonin in the WBC compartment was observed on WBC surface rather than intracellular, as observed from the partition coefficient parameters. The partition coefficient between WBC-Plasma (K_w/p_) as well as WBC-WBC-surface(K_w/wm_) were higher than one,^35^ which validate our interpretation on preferential presence of calcitonin on WBC membranes.

Calcitonin is reported to be marginally (∼ 40%) bound to plasma proteins.^1,2,32^ Hence in this study we presumed that calcitonin may similarly bind to proteins on WBC membrane as well. Most literature attribute plasma protein binding of drug to albumin due its abundance in the plasma,^24,34,35^ despite the possibility of many other plasma proteins also binding with calcitonin. While several studies have reported drugs binding to proteins on RBC membrane the literature on drugs bound to WBC membrane is relatively scant.^24,25,27,34,35^ Hence in this study to understand the types of proteins in plasma or WBC membrane having the potential to bind with calcitonin, we adopted a previously reported molecular docking approach to quantify the affinity of protein binding with both hCT and sCT. Our observation of relatively higher affinity of sCT compared to hCT towards calcitonin receptor and its signalling is consistent with previous reports.^2,5,13^ In contrast to the general assumption of higher binding of calcitonin to albumin in plasma,^2,5,13^, we observed that both hCT and sCT had weaker affinity to albumin compared to its affinity to other plasma proteins. In the plasma both hCT and sCT were observed to have superior affinity to beta-2-microglobulin’s, although the physiological significance of this is unknown. The binding of calcitonin to proteins on WBC membrane is not previously reported. On the WBC membrane hCT and sCT were observed to have relatively higher affinity towards glycoproteins (CD44), sushi domains (CD25), fibronectins (CD206) and siglecs (CD22). The binding of calcitonin with several proteins in plasma and WBC membrane at physiologically relevant affinities suggest that these interactions can favour sustained release of calcitonin to achieve its extended pharmacodynamics effects much beyond its currently known short half-life. Future studies to take advantage of these various calcitonin-protein interactions will have therapeutic merits and should be investigated.

In summary, this study reports a novel observation on sub-compartmentalization of calcitonin on WBC membranes at therapeutically relevant concentrations, which can contribute to sustained availability of calcitonin for its pharmacological effects. This study also reports several novel calcitonin-protein interactions in plasma and WBC-membrane, which merits further studies to enhance therapeutic efficacy of calcitonin.

## Acknowledgement

We thank Prof. Alan Baird for his valuable suggestion to improve this manuscript and providing calcitonin and calcitonin ELISA kit. Research support from University College Dublin-Seed funding/Output Based Research Support Scheme is acknowledged.

## References

1. Austin LA, Heath H, 3rd 1981. Calcitonin: physiology and pathophysiology. N Engl J Med 304(5):269–278.

2. Chesnut CH, 3rd, Azria M, Silverman S, Engelhardt M, Olson M, Mindeholm L 2008. Salmon calcitonin: a review of current and future therapeutic indications. Osteoporos Int 19(4):479–491.

3. Adami S, Passeri M, Ortolani S, Broggini M, Carratelli L, Caruso I, Gandolini G, Gnessi L, Laurenzi M, Lombardi A, et al. 1995. Effects of oral alendronate and intranasal salmon calcitonin on bone mass and biochemical markers of bone turnover in postmenopausal women with osteoporosis. Bone 17(4):383–390.

4. Mazzuoli GF, Passeri M, Gennari C, Minisola S, Antonelli R, Valtorta C, Palummeri E, Cervellin GF, Gonnelli S, Francini G 1986. Effects of salmon calcitonin in postmenopausal osteoporosis: a controlled double-blind clinical study. Calcif Tissue Int 38(1):3–8.

5. Kamgar-Parsi K, Hong L, Naito A, Brooks III CL, Ramamoorthy A 2017. Growth-incompetent monomers of human calcitonin lead to a noncanonical direct relationship between peptide concentration and aggregation lag time. Journal of Biological Chemistry Sep 8;292(36):14963–14976.

6. Guggi D, Krauland AH, Bernkop-Schnurch A 2003. Systemic peptide delivery via the stomach: in vivo evaluation of an oral dosage form for salmon calcitonin. J Control Release 92(1-2):125–135.

7. Lee YH, Sinko PJ 2000. Oral delivery of salmon calcitonin. Adv Drug Deliv Rev 42(3):225–238.

8. Shah RB, Palamakula A, Khan MA 2004. Cytotoxicity evaluation of enzyme inhibitors and absorption enhancers in Caco-2 cells for oral delivery of salmon calcitonin. J Pharm Sci 93(4):1070–1082.

9. Hibbins AR, Govender M, Indermun S, Kumar P, du Toit LC, Choonara YE, Pillay V 2018. In Vitro-In Vivo Evaluation of an Oral Ghost Drug Delivery Device for the Delivery of Salmon Calcitonin. J Pharm Sci 107(6):1605–1614.

10. Onishi H, Tokuyasu A 2016. Preparation and Evaluation of Enteric-Coated Chitosan Derivative-Based Microparticles Loaded with Salmon Calcitonin as an Oral Delivery System. Int J Mol Sci 17(9).

11. Carr DA, Gomez-Burgaz M, Boudes MC, Peppas NA 2010. Complexation Hydrogels for the Oral Delivery of Growth Hormone and Salmon Calcitonin. Ind Eng Chem Res 49(23):11991–11995.

12. Liu Y, Chen X, Li S, Guo Q, Xie J, Yu L, Xu X, Ding C, Li J, Ding J 2017. Calcitonin-Loaded Thermosensitive Hydrogel for Long-Term Antiosteopenia Therapy. ACS Appl Mater Interfaces 9(28):23428–23440.

13. Sato H, Tabata A, Moritani T, Morinaga T, Mizumoto T, Seto Y, Onoue S 2020. Design and Characterizations of Inhalable Poly(lactic-co-glycolic acid) Microspheres Prepared by the Fine Droplet Drying Process for a Sustained Effect of Salmon Calcitonin. Molecules 25(6).

14. Hamza A, Saramet G 2020. Actualities in Endocrine Pharmacology: Advances in the Development of Oral Formulations for Calcitonin and Semaglutide. Acta Endocrinol (Buchar) 16(3):383–387.

15. Wilkinson RJ, Vordermeier HM, Wilkinson KA, Sjolund A, Moreno C, Pasvol G, Ivanyi J 1998. Peptide-specific T cell response to Mycobacterium tuberculosis: clinical spectrum, compartmentalization, and effect of chemotherapy. J Infect Dis 178(3):760–768.

16. York-Durán MJ, Godoy-Gallardo M, Labay C, Urquhart AJ, Andresen TL, Hosta-Rigau L 2017. Recent advances in compartmentalized synthetic architectures as drug carriers, cell mimics and artificial organelles. Colloids and Surfaces. Biointerfaces Apr 1;152:199–213.

17. Overgaard K, Agnusdei D, Hansen MA, Maioli E, Christiansen C, Gennari C 1991. Dose-response bioactivity and bioavailability of salmon calcitonin in premenopausal and postmenopausal women. J Clin Endocrinol Metab 72(2):344–349.

18. Herbig ME, Weller K, Krauss U, Beck-Sickinger AG, Merkle HP, Zerbe O 2005. Membrane surface-associated helices promote lipid interactions and cellular uptake of human calcitonin-derived cell penetrating peptides. Biophys J 89(6):4056–4066.

19. Goothy SSK, Kumar AHS 2020. Network Proteins of Angiotensin-converting Enzyme 2 but Not Angiotensin-converting Enzyme 2 itself are Host Cell Receptors for SARS-Coronavirus-2 Attachment. BEMS Reports 6(1):1–5.

20. Kumar AHS 2020. Molecular Docking of Natural Compounds from Tulsi (Ocimum sanctum) and neem (Azadirachta indica) against SARS-CoV-2 Protein Targets. BEMS Reports 6(1):11–13.

21. Kumar AHS. 2021. Molecular profiling of Neprilysin expression and its interactions with SARS-CoV-2 spike proteins to develop evidence base pharmacological approaches for therapeutic intervention., ed., Online: Research Square. p 1–16.

22. Yang Z, Lasker K, Schneidman-Duhovny D, Webb B, Huang CC, Pettersen EF, Goddard TD, Meng EC, Sali A, Ferrin TE 2012. UCSF Chimera, MODELLER, and IMP: an integrated modeling system. J Struct Biol 179(3):269–278.

23. Sagar VK, Kumar AHS 2020. Efficacy of Natural Compounds from Tinospora cordifolia against SARS-CoV-2 Protease, Surface Glycoprotein and RNA Polymerase. BEMS Reports 6(1):6–8.

24. Kalaydina RV, Bajwa K, Qorri B, Decarlo A, Szewczuk MR 2018. Recent advances in “smart” delivery systems for extended drug release in cancer therapy. Int J Nanomedicine 13:4727–4745.

25. Ekladious I, Colson YL, Grinstaff MW 2019. Polymer-drug conjugate therapeutics: advances, insights and prospects. Nat Rev Drug Discov 18(4):273–294.

26. Seifert SM, Chen X, Meditz AL, Castillo-Mancilla JR, Gardner EM, Predhomme JA, Clayton C, Austin G, Palmer BE, Zheng JH, Klein B, Kerr BJ, Guida LA, Rower C, Rower JE, Kiser JJ, Bushman LR, MaWhinney S, Anderson PL 2016. Intracellular Tenofovir and Emtricitabine Anabolites in Genital, Rectal, and Blood Compartments from First Dose to Steady State. AIDS Res Hum Retroviruses 32(10-11):981–991.

27. Pardridge WM 2016. CSF, blood-brain barrier, and brain drug delivery. Expert Opin Drug Deliv 13(7):963–975.

28. Wildfeuer A, Laufen H, Zimmermann T 1994. Distribution of orally administered azithromycin in various blood compartments. International journal of clinical pharmacology and therapeutics Jul 1;32(7):356–360.

29. Treglia G, Aktolun C, Chiti A, Frangos S, Giovanella L, Hoffmann M, Iakovou I, Mihailovic J, Krause BJ, Langsteger W, Verburg FA, Luster M, Eanm, the ETC 2016. The 2015 Revised American Thyroid Association guidelines for the management of medullary thyroid carcinoma: the “evidence-based” refusal to endorse them by EANM due to the “not evidence-based” marginalization of the role of Nuclear Medicine. Eur J Nucl Med Mol Imaging 43(8):1486–1490.

30. Wells SA, Jr., Asa SL, Dralle H, Elisei R, Evans DB, Gagel RF, Lee N, Machens A, Moley JF, Pacini F, Raue F, Frank-Raue K, Robinson B, Rosenthal MS, Santoro M, Schlumberger M, Shah M, Waguespack SG, American Thyroid Association Guidelines Task Force on Medullary Thyroid C 2015. Revised American Thyroid Association guidelines for the management of medullary thyroid carcinoma. Thyroid 25(6):567–610.

31. Comba BA, Cinar A 2016. Investigation of effects of fluorosis on some minerals and hormones in sheep. Ankara Üniversitesi Veteriner Fakültesi Dergisi Sep 1;63(3):223–227.

32. Barlet JP, Godeneche D, Jourde H, Michelot J 1973. Calcitonin and iodinated hormones in sheep. Horm Metab Res 5(6):475–476.

33. Phillippo M, Lawrence CB, Bruce JB 1972. The effect of feeding on the release of calcitonin in sheep. J Endocrinol 52(1):25–30.

34. Deleu S, Kakuda TN, Spittaels K, Vercauteren JJ, Hillewaert V, Lwin A, Leopold L, Hoetelmans RMW 2018. Single-and multiple-dose pharmacokinetics and safety of pimodivir, a novel, non-nucleoside polymerase basic protein 2 subunit inhibitor of the influenza A virus polymerase complex, and interaction with oseltamivir: a Phase 1 open-label study in healthy volunteers. Br J Clin Pharmacol 84(11):2663–2672.

35. Panchagnula R, Thomas NS 2000. Biopharmaceutics and pharmacokinetics in drug research. Int J Pharm 201(2):131–150.

